# Environmental light is required for maintenance of long-term memory in *Drosophila*

**DOI:** 10.1101/592667

**Authors:** Show Inami, Shoma Sato, Shu Kondo, Hiromu Tanimoto, Toshihiro Kitamoto, Takaomi Sakai

**Author notes:** Correspondence should be addressed to Takaomi Sakai.

## Abstract

Long-term memory (LTM) is stored as functional modifications of relevant neural circuits in the brain. A large body of evidence indicates that the initial establishment of such modifications through the process known as memory consolidation requires learning-dependent transcriptional activation and *de novo* protein synthesis. However, it remains poorly understood how the consolidated memory is maintained for a long period in the brain, despite constant turnover of molecular substrates. Using the *Drosophila* courtship-conditioning assay of adult males as a memory paradigm, here, we show that in *Drosophila*, environmental light plays a critical role in LTM maintenance. LTM is impaired when flies are kept in constant darkness (DD) during the memory maintenance phase. Because light activates the brain neurons expressing the neuropeptide Pigment-dispersing factor (Pdf), we examined the possible involvement of Pdf neurons in LTM maintenance. Temporal activation of Pdf neurons compensated for the DD-dependent LTM impairment, whereas temporal knockdown of Pdf during the memory maintenance phase impaired LTM in light–dark cycles. Furthermore, we demonstrated that the transcription factor cAMP response element-binding protein (CREB) is required in the memory center, mushroom bodies (MBs), for LTM maintenance, and Pdf signaling regulates light-dependent transcription via CREB. Our results demonstrate for the first time that universally available environmental light plays a critical role in LTM maintenance by activating the evolutionarily conserved memory modulator CREB in MBs via the Pdf signaling pathway.

**Significant Statement:** Temporary memory can be consolidated into long-term memory (LTM) through *de novo* protein synthesis and functional modifications of neuronal circuits in the brain. Once established, LTM requires continual maintenance so that it is kept for an extended period against molecular turnover and cellular reorganization that may disrupt memory traces. How is LTM maintained mechanistically? Despite its critical importance, the molecular and cellular underpinnings of LTM maintenance remain elusive. This study using *Drosophila* is significant because it revealed for the first time in any organism that universally available environmental light plays an essential role in LTM maintenance. Interestingly, light does so by activating the evolutionarily conserved transcription factor cAMP response element-binding protein via peptidergic signaling.

## Introduction

A newly formed memory is initially labile, but under certain circumstances, it is consolidated into a more stable long-term memory (LTM). Previous studies using various animal species have shown that activation of specific transcription factors, such as the cAMP response element-binding protein (CREB), and the corresponding *de novo* protein synthesis are essential for memory consolidation (Yin and Tully, 1996; Lee et al., 2008; Kandel, 2012). Once consolidated, LTM requires continual maintenance for its long-term storage and subsequent recall, because memory traces gradually decay owing to molecular turnover and cellular reorganization. Similar to memory consolidation, transcriptional activation and *de novo* protein synthesis are required for LTM maintenance (Bekinschtein et al., 2007; Alberini, 2009; Majumdar et al., 2012; Fioriti et al., 2015; Hirano et al., 2016). For example, the maintenance of hippocampal long-term potentiation (LTP) and spatial memory in mice is dependent on the prion-like translational regulator CPEB3 (Fioriti et al., 2015). Similarly, LTM maintenance in *Drosophila* also requires Orb2, a *Drosophila* homolog of CPEB (Majumdar et al., 2012). Furthermore, transcriptional regulations through CREB and its coactivator CRTC play crucial roles in LTM maintenance in the *Drosophila* memory center, namely, the mushroom bodies (MBs) (Hirano et al., 2016), suggesting that memory consolidation and maintenance share some of the same molecular mechanisms (Bekinschtein et al., 2007). An obvious and important question is how transcriptional activation and the following protein synthesis are triggered during the memory maintenance phase. Unlike transcriptional and translational activations involved in memory consolidation, those associated with LTM maintenance cannot be directly controlled by stimuli that induce memory formation, because there is a significant time separation between stimulus-induced memory formation and the memory maintenance process, and LTM should be continually maintained. However, the molecular and cellular underpinnings of active LTM maintenance, which is transcription- and translation-dependent still remain elusive.

Earth’s rotation generates the daily cycle of day and night, and the rhythmic light–dark (LD) cycles have a significant impact on animal behavior and physiology. In animals, light is not only essential for acquiring information for image-forming vision in nature but also acts as a powerful modulator of brain functions such as circadian entrainment, hormone secretion, sleep–wake cycles, mood, and cognitive functions (Altimus et al., 2008; Vandewalle et al., 2009; Crocker and Sehgal, 2010; LeGates et al., 2012). Using the diurnal fruitfly *Drosophila melanogaster,* here, we found that LTM was severely impaired in flies kept in constant darkness (DD) after memory consolidation. Thus, LTM maintenance is found to be light-dependent. In *Drosophila*, light activates photoreceptors in the brain neurons expressing the Pigment-dispersing factor (Pdf), a neuropeptide, and increases their spontaneous firing rate (Sheeba et al., 2008; Fogle et al., 2011; Ni et al., 2017). Considering the physiological properties of Pdf neurons, it is possible that Pdf neurons regulate light-dependent LTM maintenance. In this study, we found that the Pdf neurons play an essential role in light-dependent LTM maintenance. Our results demonstrate for the first time that environmental light, which is available daily to all animals under normal conditions, plays a critical role in LTM maintenance by reactivating the evolutionarily conserved memory modulator CREB via Pdf signaling.

## Materials and Methods

### Fly stocks

All flies were raised on glucose-yeast-cornmeal medium in 12:12 LD cycles at 25.0 ± 0.5 °C (45–60% relative humidity). Virgin males and females were collected without anesthesia within 8 h after eclosion. The fly stocks used for this study are as follows: wild-type Canton-S (CS), 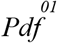 (BL26654), *Pdfr*^*5304*^(BL33068), *Pdf*-GAL4 (BL6900), c929 (BL25373), *R14F03* (BL48648), *R18F07* (BL47876), OK107 (BL854), *R61G12*-LexA (BL52685), *R41C10* (BL50121), *R55D03* (BL47656), *R19B03* (BL49830), c305a (BL30829), *nSyb*-GAL4 (BL51941), UAS-*FLP* (BL4539), UAS-*Kir2.1::eGFP* (BL6596), UAS-*TrpA1* (BL26263), UAS-*mCD8::GFP* (BL5137), UAS-*mCherry::NLS* (BL38424), UAS-*mCD8::RFP* (BL32218), UAS-*Pdf* RNAi (VDRC4380), UAS-*CrebB-B* (also known as UAS-*dCREB2-b*, BL7219), *tub*-GAL80^ts^ (BL7017), LexAop2-*FLPL* (BL55820), LexAop2-*mCD8::GFP* (BL32203), LexAop-*TrpA1* (provided by Dr. Mani Ramaswami), *hs*-*CrebB-B* (also known as *hs*-*dCREB2-b*) (Yin et al., 1994; Sakai et al., 2004), UAS-*Pdf* (provided by Dr. Taishi Yoshii), UAS-*luc* RNAi (provided by Dr. Kanae Ando), UAS>STOP>*Kir2.1::eGFP* (provided by Dr. David J. Anderson), and CRE>mCherry::STOP>*luc* (provided by Dr. Jerry C. P. Yin). All lines for behavior experiments except for 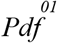, *Pdfr*^*5304*^, UAS-*luc* RNAi, LexAop2-*FLPL* were outcrossed for at least five generations to *white*^*1118*^ flies with the CS genetic background.

### Courtship-conditioning assay

The courtship-conditioning assay was carried out as previously described (Sakai et al., 2004) with some modifications. Unreceptive mated females were prepared as “trainers” one or two d before they were used for courtship-conditioning. In this conditioning, a virgin CS female and a male (3–6-d-old) were placed in an acrylic courtship chamber (15 mm in diameter × 3 mm in depth) for copulation. For LTM, a 3–5-d-old-male was placed with a mated female (4–7-d-old) in a conditioning chamber (15 mm in diameter, 5 mm in depth) containing food for 7 h either with (conditioned) or without (naïve) a single premated female (7 h conditioning). If males remated with mated females during conditioning, we discarded such males after conditioning. After 7 h conditioning, only flies showing courtship behaviors toward the mated female but not copulating successfully were transferred to a glass tube with food (12 mm in diameter × 75 mm in depth) and kept in isolation for 1 or 5 d until the test. The test was performed using a freeze-killed virgin female in a test chamber (15 mm in diameter, 3 mm in depth). All procedures in the experiments were carried out at 25 ± 1.0 °C (45–60% relative humidity) except for the temperature shift experiments. Courtship index (CI) was used for quantifying male courtship behaviors of individual flies and was calculated manually. CI is defined as the percentage of time spent in performing courtship behaviors during a given observation period (10 min). We first measured CI in conditioned and naïve males (CI_Conditioned_ and CI_Naïve_), and then mean CI_Naïve_ and mean CI_Conditioned_ were calculated. To quantify courtship memory as previously reported (Lee et al., 2017), memory index (MI) was calculated using the following formula: MI = (mean CI_Naïve_ − mean CI_Conditioned_) / mean CI_Naïve_.

### Lighting conditions in courtship-conditioning

When courtship-conditioning was performed in a dark place, conditioning chambers were placed in a temperature-regulated (25.0 ± 0.5 °C) light-tight incubator (MIR-254, Sanyo Electric Co., Ltd.). To check whether mating occurs during conditioning, we observed the flies for 10 s every 30 min by opening the incubator door. To determine whether lighting conditions affect memory maintenance, conditioned and naïve flies were kept in a light-tight incubator (MIR-254, Sanyo Electric Co., Ltd.) for one, two, or four days after 7 h conditioning.

### Temporal activation of Pdf neurons

The temperature-sensitive cation channel TrpA1 was used to activate Pdf neurons (Hamada et al., 2008). For the activation of Pdf neurons for two days during DD (the second and third days after 7 h conditioning), *Pdf*-GAL4/UAS-*TrpA1* flies were kept at 30 °C for 8 h within each subjective day [circadian time (CT) 0–8] or night (CT 12–20). However, DD-dependent LTM impairment was not improved. Since it is possible that the activation of Pdf neurons under this condition is insufficient for LTM maintenance, the flies were kept at 34 °C for 8 h within each subjective day (CT 0–8) or night (CT 12–20). UAS-*TrpA1*/+ and *Pdf*-GAL4/+ flies were used as the control.

### Temporal gene expression using TARGET system

The *tub*-GAL80^ts^ transgene used in the TARGET system (McGuire et al., 2003) encodes a ubiquitously expressed, temperature-sensitive GAL4 repressor that is active at the permissive temperature (PT, 25 °C) but not at the restrictive temperature (RT, 30 or 32 °C). By using UAS-*Pdf* RNAi, or UAS-*Pdfr* RNAi combined with the TARGET system, we knocked down *Pdf* or *Pdfr* in GAL4-positive neurons at RT (30 °C), but not at PT. In these experiments, we shifted PT to RT and vice versa during three experimental phases: 24 h before the end of conditioning, 48–72 h after conditioning, and 24 h before the initiation of the test. Furthermore, to drive the expression of a UAS-*CrebB-B* construct in MBs during a specific temporal phase, the TARGET system was also used. In *CrebB-B* experiments, we shifted PT to RT (30 °C) and vice versa during the two experimental phases: 10 h before the end of conditioning and 48–72 h after conditioning. To drive the expression of a UAS-*Kir2.1* construct in Pdf neurons during the memory maintenance phase using the TARGET system, flies were kept at RT (32 °C) for 48–72 h after conditioning. Subsequently, they were kept at PT.

### Electrical silencing of large ventral lateral clock neurons

Pdf neurons form two clusters, small lateral ventral neurons (s-LNvs) and large lateral ventral neurons (l-LNvs). To assay whether l-LNv-specific electrical silencing affects LTM, two binary gene expression systems (GAL4/UAS and LexA/LexAop) combined with Flippase (FLP/FRT) were used. The specific target gene is expressed in GAL4- and LexA-coexpressing neurons using this system,. *R61G12*-LexA and *R14F03*-GAL4 lines were used in the experiments.

### Real-time quantitative reverse transcription PCR (qRT-PCR)

A PicoPure RNA Isolation Kit (KIT0204, Thermo Fisher Scientific) was used for collecting total RNA from three whole brains in each genotype. cDNA was synthesized by the reverse transcription reaction using a QuantiTect Reverse Transcription Kit (#205311, QIAGEN). Real-time quantitative PCR was carried out using the THUNDERBIRD SYBR qPCR Mix (QPS-201, TOYOBO) and a Chromo 4 detector (CFB-3240, MJ Research). The primer sequences (custom-made by Eurofins Genomics) used for qRT-PCR were as follows: *Pdf*-Forward, 5’-ATCGGGATCTCCTCGACTGG-3’; *Pdf*-Reverse, 5’-ATGGGCCCAAGGAGTTCTCG-3’; *Pdfr*-Forward, 5’-CGTCTCATGCAGCAGATGGG-3’; *Pdfr*-Reverse, 5’-TAAGGGCGAACAGGGAGAGG-3’; *rp49*-Forward, 5’-AAGATCGTGAAGAAGCGCAC-3’; *rp49*-Reverse, 5’-TGTGCACCAGGAACTTCTTG-3’. The expression level of each mRNA was normalized by that of *rp49* mRNA. The average of the normalized mRNA expression level in control flies was calculated using data from 4–6 independent assays. The expression level of each mRNA was normalized by that of *rp49* mRNA. The average of the normalized mRNA expression level in control flies was calculated using data from 4–6 independent assays.

### Luciferase assay

To test whether light or Pdfr activation promotes CrebB activity in MBs, the Luciferase (Luc) reporter was used (Tanenhaus et al., 2012). Luc should be expressed in MBs in the combination of CRE>mCherry::STOP>*luc* reporter, MB-GAL4, and UAS-*FLP*. Three MB-GAL4 lines (*R41C10*, *R55D03*, and *R19B03*) were used in these experiments. *In vitro* Luc activity was measured using a Luciferase Assay System (E1501, Promega). Three adult male heads were collected into a 1.5 ml Eppendorf tube and homogenized in 50 μl of Glo Lysis Buffer (E266A, Promega) at Zeitgeber time (ZT) 0–2. After centrifugation, 10 μl of the resulting supernatant and 50 μl of Luciferin solution were used to analyze Luc activity. The luminescence of each sample was measured using a luminometer (GloMax 20/20, Promega) and normalized to total protein concentration using a Protein Assay Kit (#5000006, Bio-Rad). The UAS-*FLP* transgene used in this study displayed leaky expression of Luc [211–279 relative luminescence unit (RLU)/mg]. Thus, first, in the control line (UAS-*FLP*/CRE>mCherry::STOP>*luc*), the mean normalized luminescence (mean L_control_) was calculated. CrebB activity index (CAI) (RLU/mg) was defined as the difference between the “mean L_control_” and the measured luminescence in each sample in each genotype, and finally, we calculated the mean CAI. To examine whether CrebB activity increases after 7 h conditioning, we measured the CAI in each sample using conditioned or naïve flies (CAI_Conditioned_ and CAI_Naïve_), and then mean CAI_Naïve_ was calculated. Finally, we calculated the conditioning-dependent CrebB activity using the following formula: conditioning-dependent CrebB activity = (CAI_Conditioned_ − mean CAI_Naïve_) / mean CAI_Naïve_. To examine whether the induction of CrebB-B inhibits CrebB activity, we used UAS-*FLP*/*hs*-*CrebB-B*; CRE>mCherry::STOP>*luc*/*R19B03* flies. For the heat-shock treatment, 3–4-d-old male flies were grouped as 20 flies per food vial. Flies were heat-shocked at 32°C for three days. Luc activity was measured immediately after the heat-shock treatment at ZT 0–2. To examine whether the activation of Pdf neurons increases CrebB activity in the MBs, we used UAS-*FLP*/*R61G12*-LexA; CRE>mCherry::STOP>*luc* LexAop*-TrpA1*/*R41C10* flies. For the activation of Pdf neurons by TrpA1, the 3–5-d-old male flies were grouped as 20 flies per food vial. Flies were heat-shocked at 32 °C for two days during DD (the second and third days after transfer). Luc activity was measured at ZT 0–2 after flies were returned to normal LD cycle.

### Immunohistochemistry

Immunohistochemistry was performed as previously described (Shimada et al., 2016). For Pdf staining, brains were stained with a mouse anti-Pdf antibody (PDF C7-s, Developmental Studies Hybridoma Bank at the University of Iowa, 1:200) follwed by Alexa Fluor 488 anti-mouse IgG or Alexa Fluor 568 anti-mouse IgG (A11001 and A11004, Thermo Fisher Scientific) as the secondary antibodies (1:1000). For GFP staining, brains were stained with a rabbit anti-GFP antibody (A11122, Thermo Fisher Scientific, 1:200), followed by Alexa Fluor 488 anti-rabbit IgG (A11008, Thermo Fisher Scientific, 1:1000) as the secondary antibody. For Per staining, brains were stained with a rabbit anti-Per antibody (sc-33742, Santa Cruz Biotechnology, 1:250), followed by Alexa Fluor 568 anti-rabbit IgG (A11011, Thermo Fisher Scientific) as the secondary antibody (1:1000). Fluorescence signals were observed under a confocal microscope [C2^+^ (Nikon) or LSM710 (Zeiss)].

### Quantitative analysis of Pdf immunoreactivity in l-LNvs

To examine whether temporal knockdown of *Pdf* in l-LNvs by the TARGET system inhibits Pdf immunoreactivity, *Pdf*-GAL4/UAS-*Pdf* RNAi; +/*tub*-GAL80^ts^ flies were used. *Pdf*-GAL4/+; +/*tub*-GAL80^ts^ flies were used as the control. After eclosion, all flies were kept for 3–6 days at PT, and then the temperature was shifted to RT at ZT 8. After 24 h, the temperature was shifted again to PT. Subsequently, we dissected the brains for antibody staining 1 h after the temperature shift to PT (ZT 9). A confocal image stack of the brain hemisphere containing l-LNvs was Z-projected into several sequential sections. Z-sections were collected at 1 μm intervals. The signal intensity of Pdf immunoreactivity was quantified in a manually set region of interest of the cell body region in each l-LNv using the NIS elements Ar (Nikon).

### Sleep analysis

Single male flies (2–3 d old) were introduced into glass tubes (3 mm in diameter × 75 mm in length) containing fly food, and the glass tubes were set in a MB5 MultiBeam Activity Monitor (Trikinetics) to monitor the locomotor activity of individual flies. In this system, 17 independent infrared beams per glass tube were used to detect fly movement. When a fly repositions from one beam to the next, it was counted as one beam-crossing. Flies were acclimated in the glass tubes for 3 days in LD cycles at 25 °C before measuring sleep amount. Locomotor activity data were collected at 1-min intervals for 5 days and analyzed with a Microsoft Excel-based program as previously described (Kume et al., 2005). Sleep was defined as behavioral inactivity for 5 min or more (Huber et al., 2004). Total sleep amount during the day or night was analyzed as previously described (Shimada et al., 2016).

To deprive flies of sleep, a MB5 MultiBeam Activity Monitor with glass tubes each containing one naïve or conditioned fly was horizontally shaken using a shaker (NJ-022NS, Nissin). The shaker was placed inside a temperature-regulated incubator (MIR-254, Sanyo Electric Co., Ltd.). The shaking speed was set to 200 rpm. On the 2nd and 3rd days after 7 h conditioning, the MB5 MultiBeam Activity Monitor was shaken for 20 sec per 3 min during only the daytime. All experiments were carried out in 12:12 LD cycles at 25.0 ± 0.5 °C.

### Statistical analyses

All the statistical analyses were performed using IBM SPSS Statistics 22 (IBM Japan, Ltd.) or BellCurve for Excel (Social Survey Research Information Co., Ltd.), except for the comparisons of MI. In all statistical analyses except for the comparisons of MI, the Kolmogorov– Smirnov test was used to determine whether the data are normally distributed. In the statistical analysis of CI, when the data were not distributed normally, we carried out the log transformation of the data. When the basic data or transformed data are normally distributed, Student’s *t*-test was used for comparisons. When the basic data and transformed data are not distributed normally, we used the Mann–Whitney *U* test for comparisons. In the statistical analysis of MI, the permutation test with 10000 random permutations was used (H_0_, the difference between experimental and control groups is 0). The free statistical package R was used for these tests (Koemans et al., 2017). In qRT-PCR, the mean (± SEM) ratio was calculated using data from 4–6 independent assays. Since the log-transformed data were normally distributed, one-way ANOVA followed by post-hoc analysis using Scheffe’s test was used. In Luciferase assay, because the basic data were normally distributed, Student’s *t*-test was used for comparisons of two means, and one-way ANOVA followed by post-hoc analysis using Schefe’s test was carried out for multiple comparisons. In quantitative analysis of Pdf immunoreactivity, all image data were acquired under identical conditions. When the basic data or log-transformed data were distributed normally, Student’s *t*-test was used. When they were not normally distributed, we performed the Mann–Whitney *U* test. In sleep analysis, because the basic data and log-transformed data were not distributed normally, we performed nonparametric ANOVA (Kruskal–Wallis test) followed by the Steel–Dwass test for multiple comparisons.

## Results

### Light is essential for LTM maintenance

To determine whether lighting conditions affect LTM in *Drosophila*, the courtship-conditioning assay was carried out (Siegel and Hall, 1979; Sakai et al., 2004; Griffith and Ejima, 2009; Keleman et al., 2012). In this assay, males receive stressors from nonreceptive mated females (e.g., sexual rejection) to block successful mating (conditioning) (Lee et al., 2017), and memory is subsequently observed as experience-dependent courtship suppression toward virgin females. One hour conditioning generates short-term memory (STM), which persists for at least 8 h, whereas 7 h conditioning induces LTM, which persists for at least 5 days (d) (Sakai et al., 2004; Ishimoto et al., 2009; Sakai et al., 2012). The courtship activity of naïve and conditioned males was quantified using CI; subsequently, MI was calculated to quantify courtship memory (see Materials and methods). When males were conditioned for 7 h in light or darkness and the conditioned males were subsequently kept under LD cycles until the test, they showed lower courtship activity on d 5 (i.e., five days after conditioning) than naïve males, and there was no significant difference in MI between these flies [Fig. 1*A*; (1) vs. (2), Permutation test; *P* = 0.5558], indicating that conditioning in darkness has no adverse effects on LTM. However, when flies were conditioned in light and then kept in DD after the conditioning and before the test, LTM was severely impaired [Fig. 1*A*; (1) vs. (3), Permutation test, *P* = 0.0020]. DD for two consecutive days after conditioning was sufficient to impair LTM [Fig. 1*A*; (1) vs. (6), Permutation test; *P* = 0.0020], but not DD for only one day [Fig. 1*A*; (1) vs. (4), Permutation test, *P* = 0.8290; (1) vs. (5), Permutation test, *P* = 0.8902], indicating that flies cannot maintain their LTM when DD lasts more than two days. In constant light (LL), the *Drosophila* circadian clock does not work normally, and flies show arrhythmic locomotor activity (Qiu and Hardin, 1996). When flies were kept in LL after conditioning, their LTM was intact [Fig. 1*A*; (1) vs. (7), Permutation test, *P* = 0.2110], as previously reported (Sakai et al., 2004). Furthermore, the LTM of several clock mutants except for *period* (*per*) mutants is intact (Sakai et al., 2004). Thus, light input, but not the circadian clock, is necessary for LTM maintenance.

**Figure 1.**
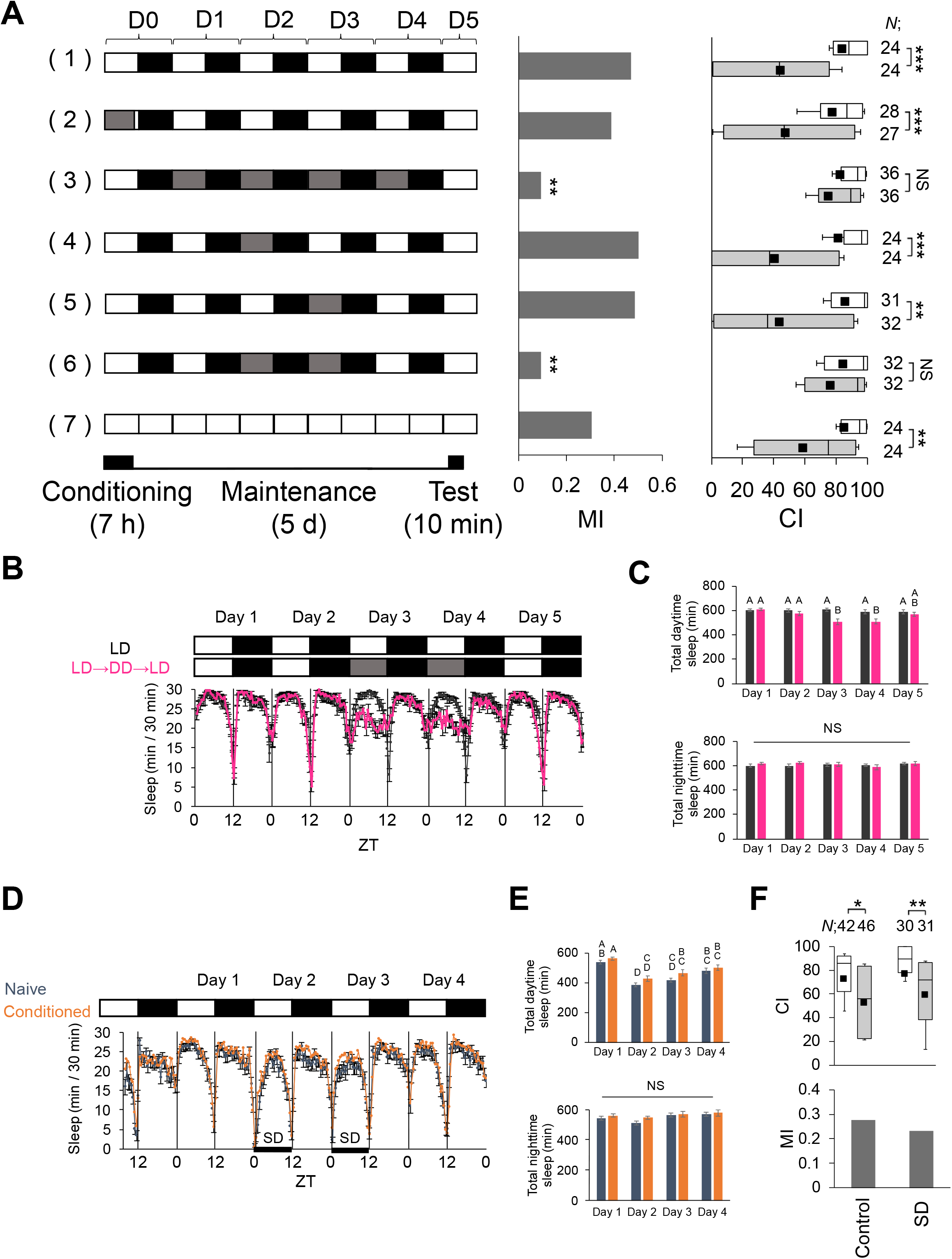
Light is essential for LTM maintenance. ***A***, LTM was measured under various lighting conditions. Wild-type males were used in the experiments. Males were tested on d 5 after 7 h conditioning. A schematic drawing of lighting conditions in courtship-conditioning is shown on the left. The white box indicates day (light) and the black box indicates night (dark). The gray box indicates that experiments were conducted in darkness during the daytime. For MI, asterisks indicate a comparison between the control (1) and test groups. ***B***–***F***, Sleep amount was measured for 5 d using wild-type males. Data are presented as mean ± SEM. ***B***, Continuous sleep amounts of control (black line) and experimental flies (magenta line). In control flies, sleep amount was measured in LD. In experimental flies, sleep amount was measured for two days in LD, which was subsequently shifted to DD for two days, and then back to LD. *N* = 31 in black line; *N* = 30 in magenta line. ***C***, Total daytime and nighttime sleep amounts in control (black bars) and experimental flies (magenta bars). NS, not significant. *N* = 31 for black bars; *N* = 30 for magenta bars. Nonparametric ANOVA (Kruskal–Wallis test) followed by post-hoc analysis using the Steel–Dwass test was carried out for multiple pairwise comparisons. Bars with the same letter indicate values that are not significantly different (*P* > 0.05). ***D***, Continuous sleep amounts in naïve (deep blue line) and conditioned flies (orange line). *N* = 30 for deep blue line; *N* = 31 for orange line. Flies were deprived of sleep during the daytime on the 2nd and 3rd days after 7 h conditioning. SD, sleep deprivation. ***E***, Total daytime and nighttime sleep amounts in naïve (deep blue bars) and conditioned flies (orange bars). Nonparametric ANOVA (Kruskal–Wallis test) followed by post-hoc analysis using the Steel–Dwass test was carried out for multiple pairwise comparisons. Bars with the same letter indicate values that are not significantly different (*P* > 0.05). *N* = 30 for deep blue bars; *N* = 31 for orange bars. ***F***, Memory on d 5 was measured using sleep-deprived flies. Control flies were kept in LD without sleep deprivation. ***A* and *F***, Box-and-whisker plots for a set of CI data show 20th, 25th, 75th, and 80th centiles. In the box-and-whisker plots, the black square in each box indicates the mean, the line in each box is drawn at the median, white boxes indicate naïve males, and gray boxes indicate conditioned males. The Mann–Whitney *U* test was used for comparisons of CI. The permutation test with 10000 random permutations was used for comparisons of MI among experimental conditions. *, *P* < 0.05; **, *P* < 0.01; ***, *P* < 0.001; NS, not significant; *N*, sample size in each box.

Sleep plays an important role in the consolidation of *Drosophila* courtship memory (Donlea et al., 2011). However, it remains unclear whether an abnormal sleep phenotype is attributable to the disturbed LTM maintenance of flies kept in DD. To examine whether light conditions affect *Drosophila* sleep (see Materials and Methods), sleep amount was measured in flies kept in LD and DD (Fig. 1*B*). The mean amount of daytime sleep of flies kept in DD was lower than that of flies kept in LD (Fig. 1*C*; total day time sleep, Kruskal–Wallis test, *H*_(9)_ = 75.113, *P* < 0.0001; total night time sleep, Kruskal–Wallis test, *H*_(9)_ = 7.534, *P* = 0.3570). However, when flies kept in LD were slightly deprived of sleep to adjust the amount of daytime sleep to the level of that in DD (Fig. 1*D* and *E*; total day time sleep, Kruskal–Wallis test, *H*_(7)_ = 75.113, *P* < 0.0001; total night time sleep, Kruskal–Wallis test, *H*_(7)_ = 7.534, *P* = 0.3570), their LTM maintenance was not attenuated (Fig. 1*F*; control vs. SD, Permutation test, *P* = 0.7204). Thus, LTM impairment induced by DD does not simply result from the reduced amount of sleep.

### Activity of Pdf neurons regulates light-dependent LTM maintenance

Since light activates Pdf neurons, it is possible that the activation of Pdf neurons restores LTM impairment induced by DD. Thus, we examined whether the temporal activation of Pdf neurons induced by the temperature-sensitive cation channel TrpA1 can compensate for the DD-dependent LTM impairment (Fig. 2*A*–*D*). When Pdf neurons were activated in *Pdf*-GAL4/UAS-*TrpA1* flies on each subjective day or night in DD for 2 d after conditioning, LTM was maintained for 5 d in *Pdf*-GAL4/UAS-*TrpA1* flies (Fig. 2*C*, *D*; Permutation test; *C*, UAS control vs. F_1_, *P* = 0.0090, GAL4 control vs. F1, *P* = 0.0038; *D,* UAS control vs. F_1_, *P* = 0.0074, GAL4 control vs. F_1_, *P* = 0.0122). Under the same temperature-shift conditions, GAL4 and UAS control flies still showed LTM impairment (Fig. 2*C*, *D*). Thus, this finding indicates that the activation of Pdf neurons during either a subjective day or night is sufficient to rescue the LTM impairment. Consistently, the electrical silencing of Pdf neurons during the memory maintenance phase attenuated LTM in LD (Fig. 2*E*, *F*; Permutation test; *E*, *P* = 0.9492; *F*, *P* = 0.0014). Taken together, these results indicate that the activity of Pdf neurons regulates light-dependent LTM maintenance.

**Figure 2.**
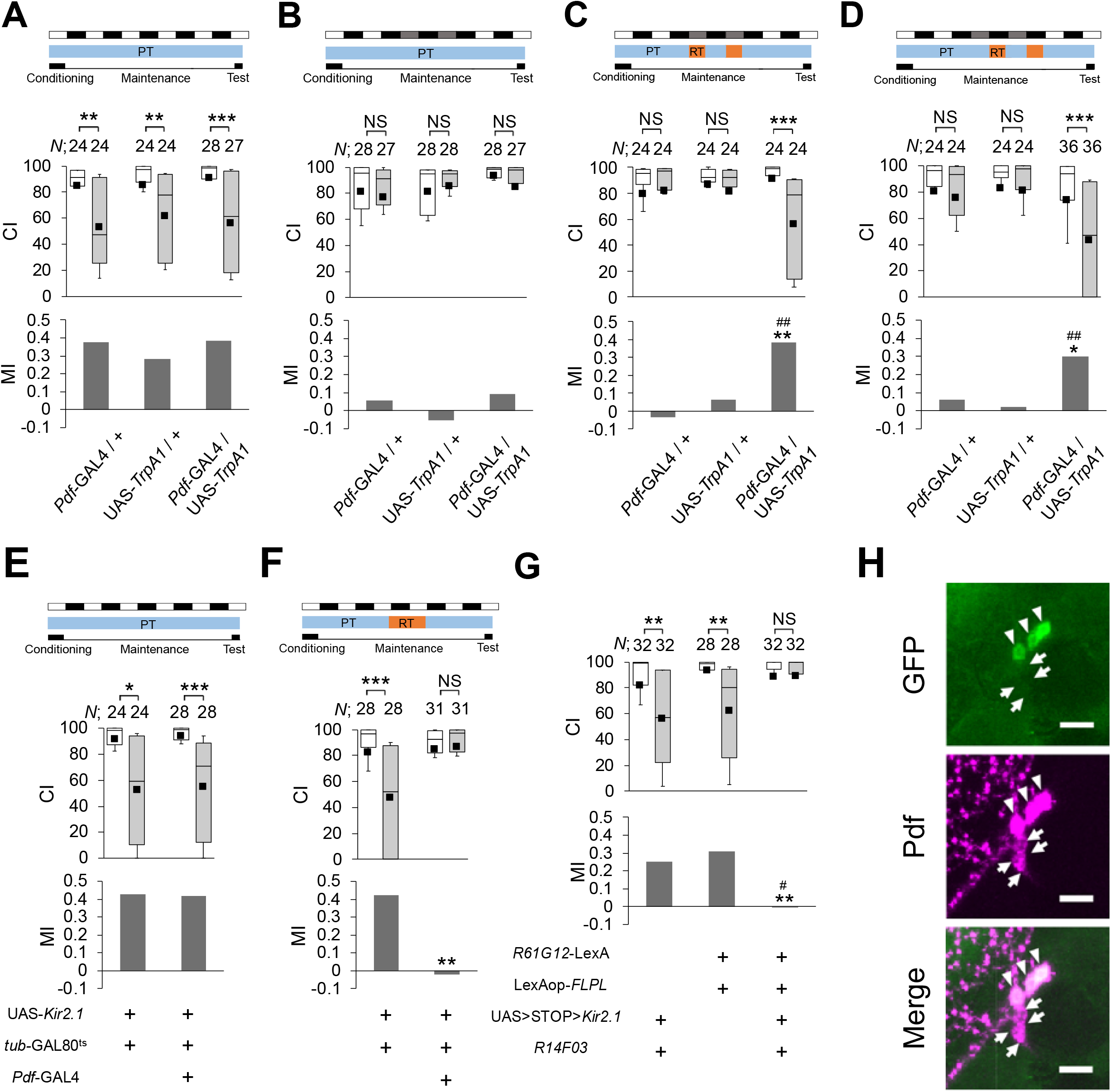
Activity of Pdf neurons is required for LTM maintenance. ***A***–***D*** Temporal activation of Pdf neurons compensates for DD-dependent LTM impairment. *Pdf*-GAL4/+, UAS-*TrpA1*/+, and *Pdf*-GAL4/UAS-*TrpA1* males were used. For MI, asterisks indicate a comparison between F_1_ (GAL4/UAS) and GAL4 control flies, and hash marks indicate a statistical comparison between F_1_ and UAS control flies. ***A***, All experiments were performed at PT. ***B***, On the 2nd and 3rd days after 7 h conditioning, flies were kept in DD. ***C***, On the 2nd and 3rd days after 7 h conditioning, flies were kept at 34°C during the period between CT 0 and CT 8. ***D***, On the 2nd and 3rd days after 7 h conditioning, flies were kept at 34°C during the period between CT 12 and CT 20. ***E* and *F***, UAS-*Kir2.1*/*Pdf*-GAL4; *tub*-GAL80^ts^/+ flies were used. UAS-*Kir2.1*/*tub*-GAL80^ts^ flies were used as the control. ***E***, All experiments were carried out at PT (25°C). ***F***, Flies were kept at RT (32°C) for 48–72 h after 7 h conditioning. ***G***, l-LNv-specific silencing impairs LTM. *R61G12*-LexA/LexAop-*FLPL*; UAS>STOP>*Kir2.1*/*R14F03* flies were used. For MI, asterisks indicate a comparison between l-LNv-silenced flies and the LexA/LexAop control (*R61G12*-LexA/LexAop-*FLPL*), and the hash mark indicates a comparison between l-LNv-silenced flies and the GAL4/UAS control (UAS>STOP>*Kir2.1*/*R14F03*). The permutation test was used (LexA/LexAop control vs. l-LNv-silenced flies, *P* = 0.0011; GAL4/UAS control vs. l-LNv-silenced flies, *P* = 0.0113). ***A***–***G***, The Mann–Whitney *U* test was used for comparisons of CI. The permutation test with 10000 random permutations was used for comparisons of MI. *, *P* < 0.05; **, *P* < 0.01; ***, *P* < 0.001; #, *P* < 0.05; ##, *P* < 0.01; NS, not significant; *N*, sample size in each box. ***H***, Confocal section images at the level of Pdf neurons of the adult brain. Scale bars represent 10 μm. Triangles, l-LNvs; arrows, s-LNvs.

### Pdf expression is critical for LTM maintenance in LD

We next sought to determine the temporal requirement of Pdf for LTM maintenance. For this purpose, we performed temporal knockdown of *Pdf* in Pdf neurons using the TARGET system (McGuire et al., 2003) and RNA interference (RNAi) technology. To demonstrate that the Pdf signaling pathway regulates light-driven LTM maintenance, we performed temporal knockdown of *Pdf* in Pdf neurons (Fig. 3*A*–*D*). The effectiveness of *Pdf* RNAi was confirmed by qRT-PCR (Fig. 4*A*; One-way ANOVA, *F*_(2, 12)_ = 6.954, *P* = 0.0099; Scheffe’s multiple comparisons, GAL4 control vs. F_1_, *P* = 0.0193, UAS control vs. F_1_, *P* = 0.0291) and immunostaining using an anti-Pdf antibody (Fig. 4 *B, C*; Student’s *t*-test in *B*, *t*_(49)_ = −1.801, *P* = 0.0778; Mann–Whitney *U* test in *C*, *U* = 365, *P* < 0.0001). To knockdown Pdf during the memory consolidation, maintenance or test phase, the temperature was raised to 30 °C for 24 h during the three experimental periods (Fig. 3*B*–*D*): starting at 24 h before the end of conditioning, 48–72 h after conditioning (memory maintenance phase), and 24 h before the test initiation. When flies were kept for 24 h before the test initiation at the restrictive temperature (RT), Pdf should remain suppressed during the 10 min test. LTM was impaired only when *Pdf* was knocked down during the memory maintenance phase (Fig. 3 *A*–*D*; Permutation test; *A*, *P* = 0.3274; *B*, *P* = 0.1904; *C*, *P* = 0.0034; *D*, *P* = 0.1290), indicating that *Pdf* expression is required for LTM maintenance. Since memory on d 1 remianed intact in 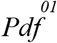 null mutant flies after 7 h conditioning (Fig. 3*E*; Permutation test, *P* = 0.9646), Pdf was found to be dispensable for memory consolidation. Unlike the memory on d 1, 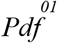 mutant flies showed memory impairment on d 5, which was rescued by *Pdf* expression (Fig. 4*F*; Permutation test, *P* = 0.0020). This finding also supports the idea that Pdf is required for keeping LTM more than one day.

**Figure 3.**
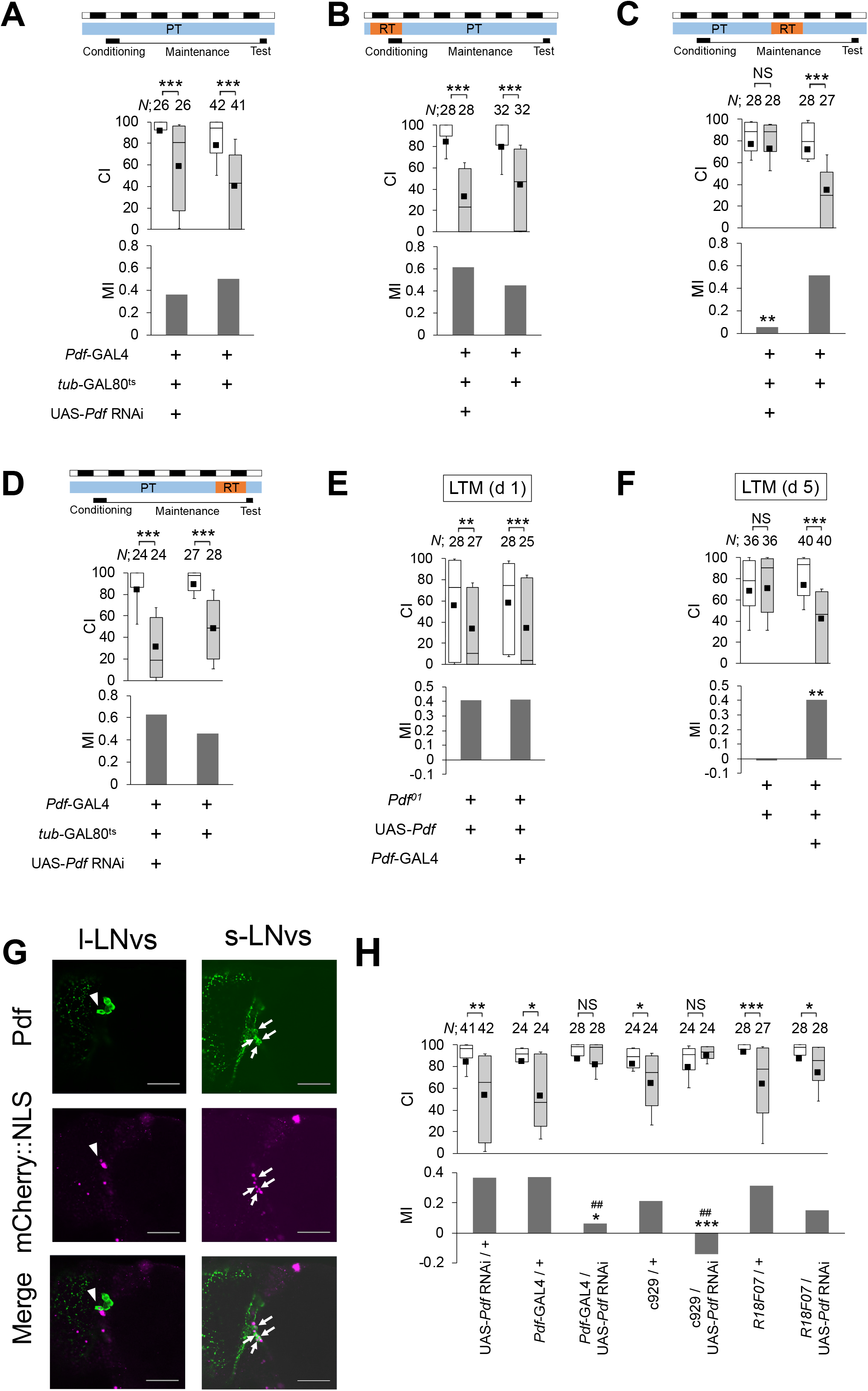
Pdf expression is essential for LTM maintenance. ***A–F* and *H***, Males were tested on d 1 or d 5 after 7 h conditioning. In the box-and-whisker plots, the white boxes indicate naïve males and the gray boxes indicate conditioned males. The Mann–Whitney *U* test was used for comparisons of CI. The permutation test with 10000 random permutations was used for comparisons of MI. *, *P* < 0.05; **, *P* < 0.01; ***, *P* < 0.001; ##, *P* < 0.01; NS, not significant; *N*, sample size in each box. ***A***, All experiments were performed at PT (25°C). ***B***, Flies were kept at RT for 24 h before the end of conditioning. ***C***, Flies were kept at RT for 48–72 h after 7 h conditioning. ***D***, Flies were kept at RT for 24 h before the test. ***A–D***, memory on d 5 in *Pdf*-GAL4 / UAS-*Pdf* RNAi; *tub*-GAL80^ts^/+ flies and control flies (*Pdf*-GAL4 /+; *tub*-GAL80^ts^/+). ***E***, Memory on d 1 in UAS-*Pdf* / *Pdf*-GAL4; 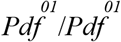 flies and control flies (UAS-*Pdf*/+; 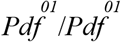). ***F***, Memory on d 5 in UAS-*Pdf* / *Pdf*-GAL4; 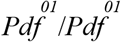 flies and control flies. ***G***, Confocal section images at Pdf neuron level of the adult brain. Triangles, l-LNvs; arrows, s-LNvs. Scale bars represent 50 μm. UAS*-mCherry::NLS*/*R18F07* flies were used. Magenta, mCherry::NLS; green, Pdf. ***H***, Knockdown of *Pdf* in l-LNvs and/or s-LNvs using three GAL4 drivers (*Pdf*-GAL4, c929, and *R18F07*). For MI, asterisks indicate a comparison between F_1_ and GAL4 control flies, and hash marks indicate a statistical comparison between F_1_ and UAS control flies. The permutation test was used (UAS control vs. c929/UAS-*Pdf* RNAi, *P* = 0.0028; GAL4 control vs. c929/UAS-*Pdf* RNAi, *P* < 0.0001; UAS control vs. *R18F07*/UAS-*Pdf* RNAi, *P* = 0.0588; GAL4 control vs. *R18F07*/UAS-*Pdf* RNAi, *P* = 0.1536).

**Figure 4.**
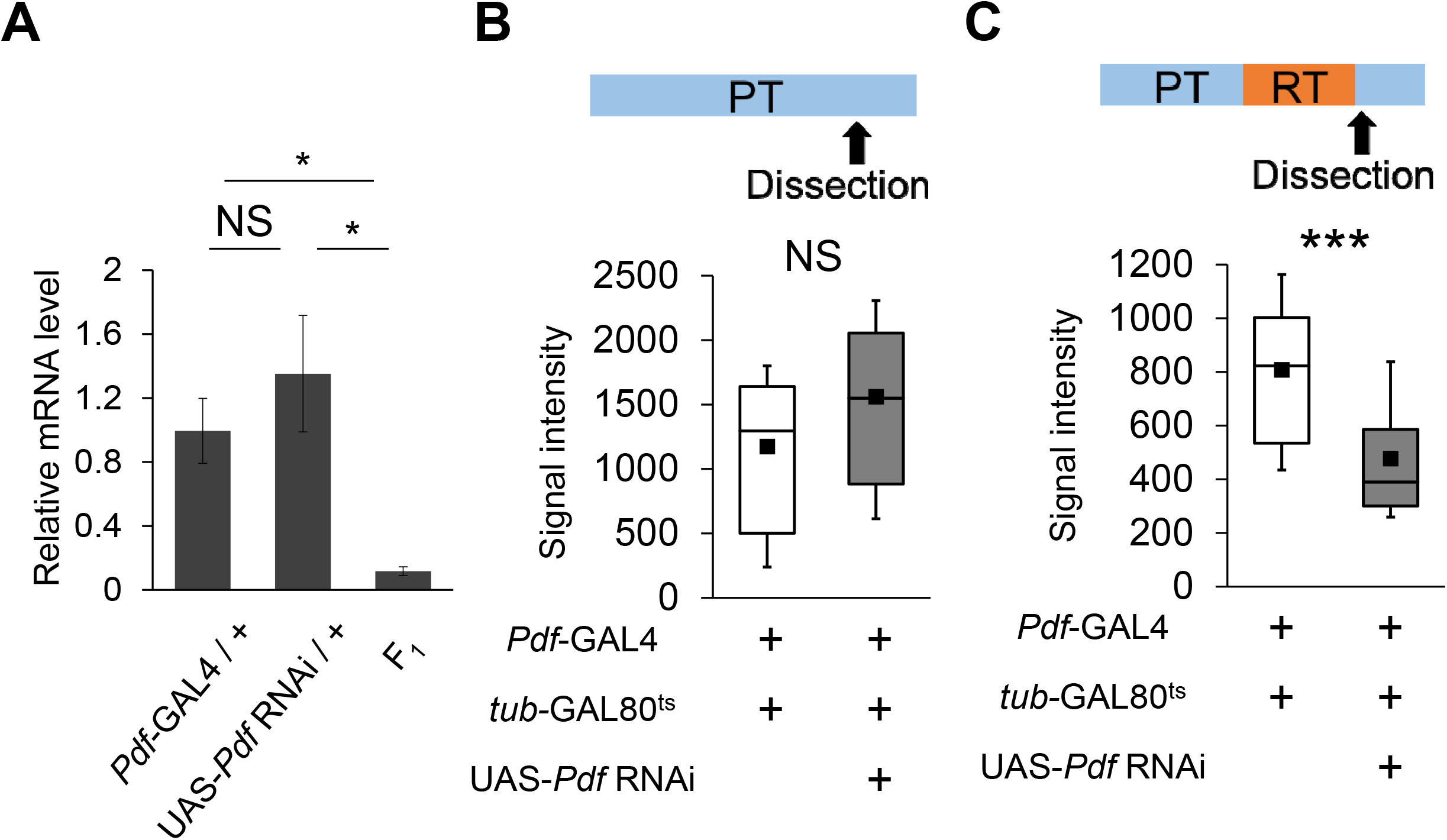
Real-time qRT-PCR analysis and quantitative analysis of Pdf immunoreactivity. **A,** Analysis of *Pdf* mRNA expression level. *Pdf*-GAL4 was used for the induction of *Pdf* RNAi. Mean ± SEM was calculated from five to six replicates. One-way ANOVA followed by post-hoc analysis using Scheffe’s test for multiple pairwise comparisons was used. *, *P* < 0.05; NS, not significant. ***B and C***, Quantitative analysis of Pdf immunoreactivity in l-LNvs. Adult brains were dissected at ZT 9. In the box- and-whisker plots, white boxes indicate *Pdf*-GAL4/+; *tub*-GAL80^ts^/+ flies and gray boxes indicate *Pdf*-GAL4/ UAS-*Pdf* RNAi; *tub*-GAL80^ts^/+ flies. ***B***, All experiments were carried out at PT (25°C). *N* = 26 for white box; *N* = 27 for gray box. Student’s *t*-test was used for statistical analysis. NS, not significant. ***C***, Flies were kept at RT (30°C) for 24 h. *N* = 50 for white box; *N* = 38 for gray box. The Mann–Whitney *U* test was used for statistical analysis. ***, *P* < 0.001.

We next used two additional GAL4 drivers, c929 and *R18F07*, to further investigate neuronal cell types involved in Pdf-mediated LTM maintenance. c929 drives the expression of GAL4 in peptidergic neurons including l-LNvs but not s-LNvs (Taghert et al., 2001; Shimada et al., 2016), and *R18F07* drives GAL4 expression in all s-LNvs and only weakly in one of the l-LNvs (Fig. 3*G*). *Pdf* knockdown in c929-positive neurons impaired LTM, but not that in *R18F07*-positive neurons (Fig. 3*H*; probability in each permutation test is shown in the figure legend). In addition, we examined whether l-LNv-specific electrical silencing impairs LTM. We confirmed that *Kir2.1::eGFP* is expressed in all l-LNvs but not in s-LNvs in *R61G12*-LexA/LexAop-*FLPL*; UAS>STOP>*Kir2.1::eGFP/R14F03* flies (Fig. 2H); moreover, l-LNv-specific electrical silencing also impaired LTM (Fig. 2*G*; the probability in each permutation test is shown in the figure legend). Taken together, it is most likely that Pdf expression in l-LNvs is essential for LTM maintenance.

### Pdfr is essential for light-dependent CrebB activity in MB α/β neurons

A *Drosophila* homolog of CREB (CrebB) is required for the consolidation and maintenance of olfactory memory (Yin and Tully, 1996; Hirano et al., 2016). Our previous studies demonstrated that the consolidation of courtship memory is also regulated by CrebB (Sakai et al., 2004; Ishimoto et al., 2009). Thus, using a repressor isoform of CrebB (CrebB-B; also known as dCREB2-b), we first examined whether CrebB is also required for the maintenance of courtship memory (Fig. 5*A*–*C*). LTM was attenuated when CrebB-B was expressed in MB α/β neurons (Fig. 5*D*) during conditioning (Fig. 5*B*; Permutation test; *P* < 0.0001) or the memory maintenance phase (Fig. 5*C*; Permutation test; *P* = 0.0054), indicating that CrebB in MB α/β neurons is required for both the consolidation and maintenance of LTM. On the other hand, CrebB in MB γ neurons (Fig. 5*E*) during conditioning, but not during the maintenance phase, attenuated LTM (Fig. 5*B, C*; Permutation test in *B*; *P* = 0.0022; Permutation test in *C*, *P* = 0.37), suggesting that CrebB in MB γ neurons is required only for memory consolidation. Unlike the MB α/β and γ neurons, MB α’/β’ neurons (Fig. 5*F*) had no effect on CrebB-dependent LTM (Fig. 5*A*–*C*; Permutation test in *B*; *P* = 0.2958; Permutation test in *C*; *P* = 0.5666).

**Figure 5.**
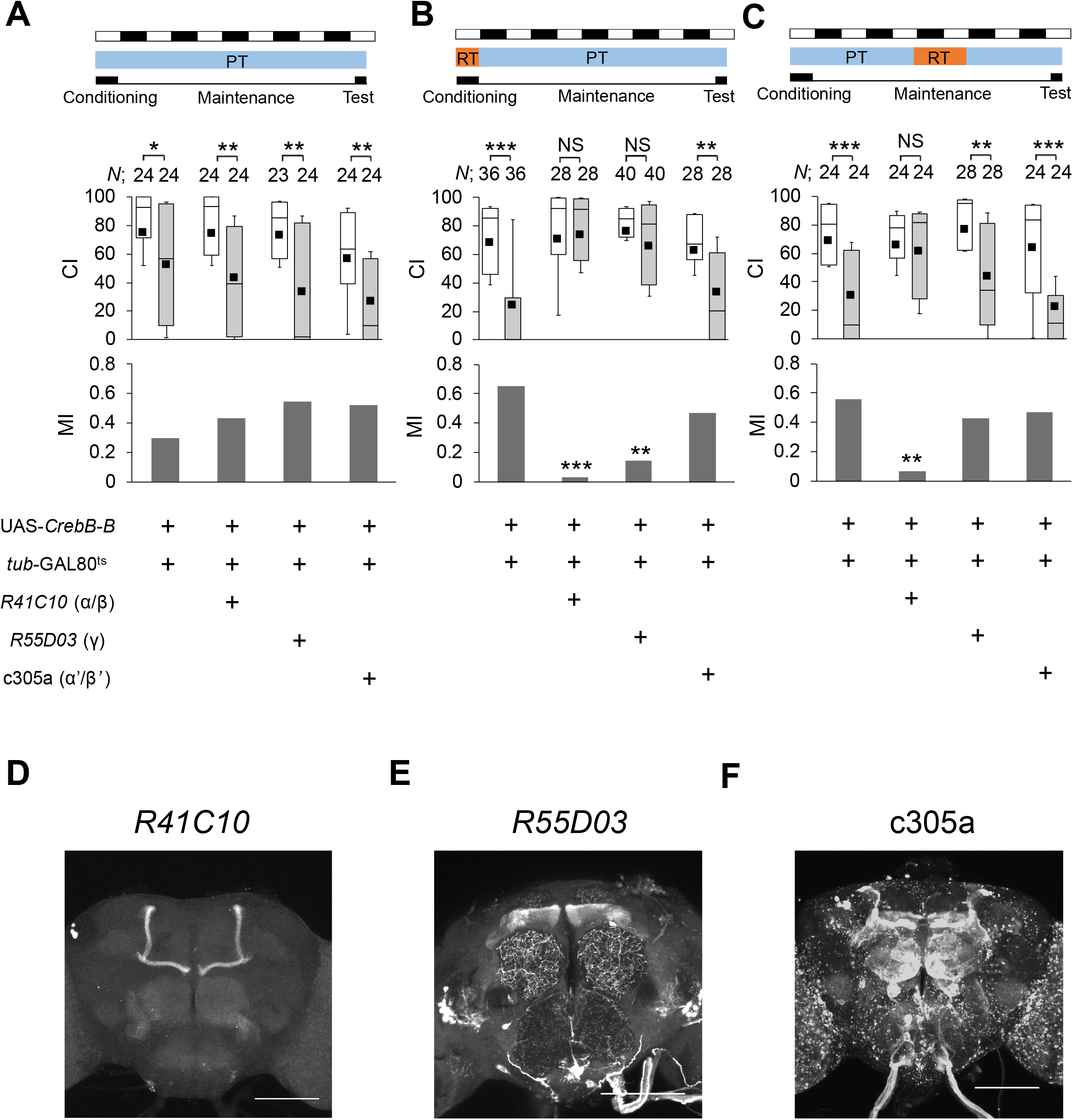
Temporal expression of CrebB repressor in MB neurons. ***A–C***, The Mann–Whitney *U* test was used for comparisons of CI. The permutation test with 10000 random permutations was used for comparisons of MI. For MI, asterisks indicate a comparison between the UAS control and test groups. *, *P* < 0.05; **, *P* < 0.01; ***, *P* < 0.001; NS, not significant; *N*, sample size in each box. All flies were raised at 25°C. The temperature was shifted during two experimental periods. ***A***, All experiments were performed at PT (25°C). ***B***, Flies were kept at RT (30°C) for 10 h before the end of conditioning. ***C***, Flies were kept at RT for 48–72 h after 7 h conditioning. ***D–F***, Stacked confocal images of the adult brain. *R41C10*/ UAS-*mCD8::GFP* **(D)**, *R55D03*/UAS-*mCD8::GFP* **(E)**, and c305a/UAS-*mCD8::RFP* **(F)**flies were used. Scale bars represent 100 μm.

We next examined whether light activates CrebB transcription in MBs. To estimate CrebB activity, we used the CRE>mCherry::STOP>*luciferase* (*luc*) reporter, which is a *luc*-based reporter gene under the control of CrebB-binding sites (CRE) (Tanenhaus et al., 2012). Using this reporter construct, in which the mCherry::STOP sequence is flipped out in the presence of the GAL4-induced recombinase FLP, we can measure the MB-specific transcriptional activity of CrebB with MB-GAL4. The induction of a CrebB repressor driven by a heat-shock promoter severely attenuated the CRE-dependent Luc activity in MBs (Fig. 6*A*; Mann–Whitney *U* test, *U* = 90, *P* < 0.0001), demonstrating that this reporter can reliably be used as an indicator of CrebB activity. When naïve flies were kept in LD, robust CrebB activity was detected in MB α/β neurons, but not in MB γ neurons [Fig. 6*B*; (1) and (2); Statistic values are shown in the figure legend]. When they were kept in DD for 1 d, the CrebB activity in MB α/β neurons decreased by approximately 50% compared with that in control flies [Fig. 6*B*; (3)]. On the second day of DD, it decreased further [Fig. 6*B*; (4)], indicating that the CrebB activity in MB α/β neurons is light-dependent. Furthermore, we examined whether the Pdf receptor (Pdfr) regulates the light-dependent CrebB activity. As was observed in 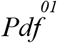 flies, *Pdfr* null mutant (*Pdfr*^*5304*^) flies also showed severe LTM impairment [Fig. 6*C*; (1) vs (5); Permutation test, *P* = 0.0034]. In LD, CrebB activity was severely attenuated in MB α/β neurons with the *Pdfr* null mutant background (Fig. 6*B*). Furthermore, in DD, the activation of Pdf neurons by TrpA1 increased the CrebB activity in MB α/β neurons (Fig. 6*D*; Mann–Whitney *U* test, *U* = 35, *P* = 0.0043). Thus, the Pdf/Pdfr signaling pathway is essential for light-driven CrebB activity in MB α/β neurons.

**Figure 6.**
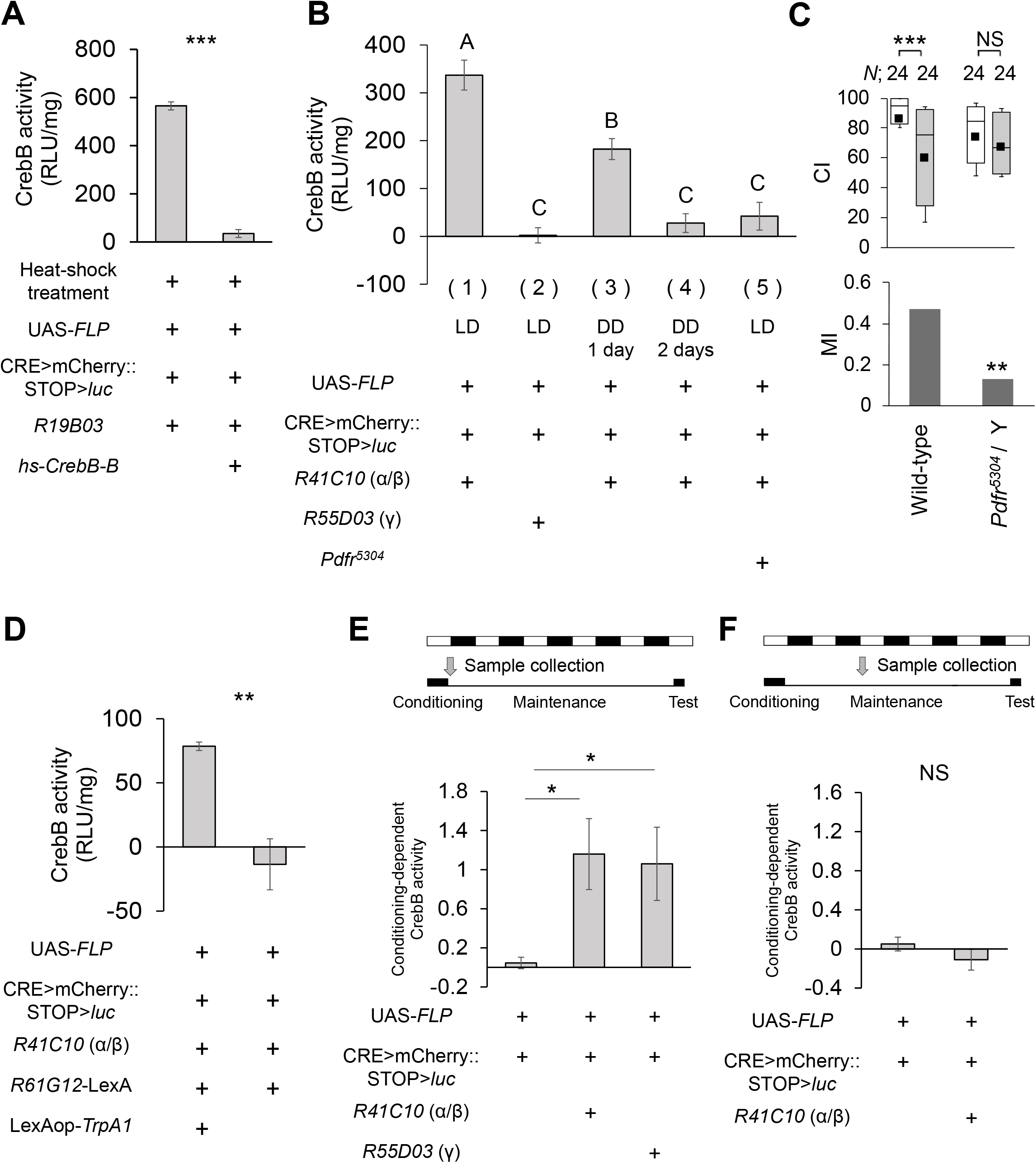
Light-dependent transcriptional activity of CrebB in MB α/β neurons. ***A***, Using UAS-*FLP*/*hs*-*CrebB-B*; CRE>mCherry::STOP>*luc*/*R19B03* and UAS-*FLP*/+; CRE>mCherry::STOP>*luc*/*R19B03* flies, we examined whether the induction of the CrebB repressor CrebB-B inhibits CrebB activity. *N* = 9–10 for each bar. In heat-shock treatment, males (3–4 d old) were kept at 32°C for three days. Luc activity was measured immediately after the heat-shock treatment. Error bars indicate SEM. Student’s *t*-test was used. ***, *P* < 0.001. ***B***, The CrebB activity in MB α/β or γ neurons was measured in LD or DD using MB α/β-GAL4 (*R41C10*) or MB γ-GAL4 (*R55D03*). Samples were prepared between ZT 0 and ZT 2. *N*= 8–13 in each bar. Error bars indicate SEM. One-way ANOVA followed by post-hoc analysis using Scheffe’s test was used (One-way ANOVA, *F*_(4, 50)_ = 31.697, *P* < 0.0001; Scheffe’s multiple comparisons, (1) vs. (2), *P* < 0.0001, (1) vs. (3), *P* = 0.0169, (1) vs. (4), *P* < 0.0001, (1) vs. (5), *P* < 0.0001, (2) vs. (3), *P* < 0.0001, (2) vs. (4), *P* = 0.9427, (2) vs. (5), *P* = 0.5709, (3) vs. (4), *P* = 0.0039, (3) vs. (5), *P* = 0.0075, (4) vs. (5), *P* = 0.9820). Bars with the same letter indicate values that are not significantly different (*P* > 0.05). ***C***, Memory on d 5 in wild-type and *Pdfr*^*5304*^ flies. The Mann–Whitney *U* test was used for comparisons of CI. The permutation test with 10000 random permutations was used for comparisons of MI. For MI, asterisks indicate a comparison between the UAS control and test groups. **, *P* < 0.01; ***, *P* < 0.001; NS, not significant; *N*, sample size in each box. ***D***, UAS-*FLP*/*R61G12*-LexA; CRE>mCherry::STOP>*luc* LexAop*-TrpA1*/*R41C10* and UAS-*FLP*/*R61G12*-LexA; CRE>mCherry::STOP>*luc* /*R41C10* flies were used. *N* = 5–6 for each bar. For activation of Pdf neurons by TrpA1, males (3–5 d old) were kept at 32°C during two days of DD. Error bars indicate SEM. Student’s *t*-test was used. **, *P* < 0.01. ***E* and *F***, The conditioning-dependent CrebB activity in MB α/β or γ neurons was measured immediately after 7 h conditioning **(E)**and on the 2nd day after 7 h conditioning **(F)**. Student’s *t*-test was used. *, *P* < 0.05; NS, not significant. *N*= 6–9 in each bar.

Next, the effect of 7 h conditioning on CrebB activity was examined. The CrebB activity in MB α/β and γ neurons increased immediately after 7 h conditioning (Fig. 6*E*; α/β neurons, Student’s *t*-test, *t*_(17)_ = −3.1965, *P* = 0.0053; γ neurons, Mann–Whitney *U* test, *U* = 78, *P* = 0.0058). This finding is consistent with a previous report (Ishimoto et al., 2009). However, no conditioning-dependent increase in CrebB activity was observed during the memory maintenance phase (Fig. 6*F*; Student’s *t*-test, *t*_(13)_ = 1.1219, *P* = 0.2822).

## Discussion

In nocturnal mice, an ultradian LD cycle condition induces spatial learning defects, although such an aberrant light condition does not impair the molecular clock or sleep amount (LeGates et al., 2012). The ultradian LD cycle also attenuates hippocampal long-term potentiation (Fernandez et al., 2018). In contrast, a short pulse of white light during the night before learning enhances long-lasting fear memory through the activation of hippocampal p21-activated kinase 1 (Shan et al., 2015). In humans, who are naturally diurnal, prior exposure to orange light promotes working memory, indicating that light also modulates human cognitive brain functions (Chellappa et al., 2014). Thus, regardless of the nocturnal or diurnal nature of animals, lighting conditions can positively or negatively modify the acquisition or consolidation of memories (Cajochen et al., 2011; LeGates et al., 2012; Chellappa et al., 2014; Shan et al., 2015). However, to the best of our knowledge, it has never been studied whether environmental light affects LTM maintenance. In this study using *Drosophila*, we demonstrated for the first time that environmental light is an essential factor for the appropriate maintenance of LTM.

The *Drosophila* Pdf neuropeptide regulates various biological phenomena such as circadian behavioral rhythms, light-driven arousal, geotactic behavior, rival-induced prolonged mating, and sex pheromone biosynthesis (Renn et al., 1999; Helfrich-Forster et al., 2000; Mertens et al., 2005; Kim et al., 2013; Krupp et al., 2013). In the present study, we found that *Drosophila* has a light-driven memory maintenance system, and that the Pdf signaling pathway regulates this system. Pdf expression was essential for LTM maintenance in LD (Fig. 3). LTM was impaired when flies were kept in DD for 2 d after 7 h conditioning (Fig. 1), and the activation of Pdf neurons was sufficient to rescue the LTM impairment induced by DD (Fig. 2). Moreover, the electrical silencing of Pdf neurons impaired LTM in LD (Fig. 2). Considering that light activates Pdf neurons (Sheeba et al., 2008; Fogle et al., 2011; Ni et al., 2017), it is most likely that light-inducible Pdf released from Pdf neurons regulates LTM maintenance. We further confirmed that Pdfr expression is necessary for the light-driven transcription through CrebB, which is essential for LTM maintenance (Fig. 5, 6). Taken together, our study shows that this light-dependent transcription system in MB α/β neurons via the Pdf signaling pathway regulates LTM maintenance in *Drosophila*.

*Drosophila* has three light-sensing organs: the compound eyes, ocelli, and Hofbauer–Buchner (H–B) eyelets. The compound eyes play key roles in light entrainment of the circadian clock (Helfrich-Forster et al., 2002), and l-LNvs receive visual information via the compound eyes (Muraro and Ceriani, 2015). The H–B eyelets are also important for circadian photoreception, and axons of H–B eyelet photoreceptors project to the circadian pacemaker neurons including l-LNvs (Li et al., 2018). In addition, light directly activates Pdf neurons through brain photoreceptors, Rhodopsin 7, and Cryptochrome (Sheeba et al., 2008; Fogle et al., 2011; Ni et al., 2017). Although light input pathways associated with light-dependent LTM maintenance remain unclear, the photoactivation of light-sensing organs and/or Pdf neurons will trigger light-dependent LTM maintenance in *Drosophila*.

The targeted expression of a CrebB repressor in MB α/β neurons during 7 h conditioning impaired LTM, as was observed in that in MB γ neurons (Fig. 5), indicating that CrebB-dependent transcription in both MB α/β and γ neurons is necessary for memory consolidation. In addition, the targeted expression of a CrebB repressor in MB α/β neurons during the memory maintenance phase also impaired LTM (Fig. 5), whereas, that in MB γ neurons did not (Fig. 5). It has been reported that MB γ neurons are necessary for 1 d memory of courtship-conditioning (Kruttner et al., 2015). Thus, consolidated courtship memory may last in MB γ neurons only for 1–2 d at most, whereas longer-lasting LTM seems to be established and stored in MB α/β neurons. In contrast to the MB γ neurons, the light-dependent activation of CrebB transcription was evident in MB α/β neurons, indicating that 5 d memory is maintained in MB α/β neurons through the Pdf/Pdfr/CrebB pathway in LD. Thus, long-lasting LTM (> 1 d memory) seems to be established and stored in MB α/β neurons.

When flies were kept in DD for two consecutive days, LTM was impaired (Fig. 1) and the CrebB activity in MB α/β neurons was severely attenuated (Fig. 6). However, DD for only 1 day was not sufficient to impair LTM (Fig. 1) but it induced a 50% reduction in CrebB activity (Fig. 6). Thus, when the CrebB activity in MB α/β neurons is severely attenuated, LTM maintenance may break down. Can Pdfr be the only factor that regulates the CrebB activity in MB α/β neurons? Pdf/Pdfr signaling should play a role in the regulation of the CrebB activity in MB α/β neurons because a *Pdfr* null mutation severely attenuated the CrebB activity (Fig. 6). However, it remains possible that signaling pathways other than the Pdf/Pdfr signaling pathway also contribute to the CrebB activity in MB α/β neurons.

In contrast to LTM maintenance, LTM formation occurred in darkness (Fig. 1). This finding is consistent with a previous report that STM after 1 h conditioning under a dim red light, which blocks visual input, remains intact (Joiner and Griffith, 1997). In this study, we found that CrebB activity during conditioning was necessary for memory consolidation (Fig. 5). Considering that flies were able to establish LTM when they were conditioned in darkness (Fig. 1), it is likely that light-independent CrebB transcription in MBs plays an important role in memory consolidation. Unlike memory consolidation, LTM maintenance requires a light-dependent transcription system in MB α/β neurons via the Pdf signaling pathway. In naïve males, however, light can also increase the CrebB transcription activity in MB α/β neurons (Fig. 6). In naïve males, it is most unlikely that this light-driven transcription system provides proteins that are required for LTM maintenance. Thus, it may play a role in innate brain functions other than LTM maintenance. Assuming that this hypothesis is true, how are proteins that are required for LTM maintenance provided in only conditioned males? Although it remains unclear, repetitive exposure to stressors using mated females during courtship-conditioning may trigger the change in the target genes of CrebB in the light-driven transcription system, and such a system in MB α/β neurons may provide gene products that are required for maintaining consolidated LTM in a courtship-conditioning-dependent manner.

In nature, animals learn much from their experience throughout the day. Through their experience, LTM should be formed and maintained for a long period. If environmental light, which is available daily to all animals in nature, can be used for transcriptional activation in the brain, such a light-driven transcription system is considered reasonable and effective for continually providing *de novo* protein synthesis required for LTM maintenance. When diurnal Nile grass rats were housed under dim LD cycles, 24 h spatial memory is impaired, and this lighting condition inhibits the expression of brain-derived neurotrophic factors and the dendritic spine density in the hippocampus (Soler et al., 2018). Although it is as yet unclarified whether the rapid forgetting in the Nile grass rats under dim LD cycles results from the inhibition of *de novo* protein synthesis required for LTM maintenance, light-dependent *de novo* protein synthesis in the memory center may be conserved in many animal species. As is observed in *Drosophila*, in mammals, repeated exposure to stressors also induces a long-lasting reduction of male sexual motivation (Hawley et al., 2011; Hawley et al., 2013). It is thus interesting to examine whether environmental light and the evolutionarily conserved memory modulator CREB also play critical roles in the maintenance of such depressed sexual motivation in mammals including humans.

## Acknowledgments

We thank Yuto Kurata and Yushun Sato for technical assistance. We also thank Toshiro Aigaki for helpful discussions. This work was supported by JSPS KAKENHI Grants (Numbers 18H04887 and 16H04816 to T.S., 17H01378, 17H05545, 16H01496, and 26250001 to H.T., and 15J06303 to S.I.).

